## Supplementary figures and images for "Lipopolysaccharide-mediated resistance to host antimicrobial peptides and hemocyte-derived reactive-oxygen species are the major *Providencia alcalifaciens* virulence factors in *Drosophila melanogaster*"

### Supplemental figure 1

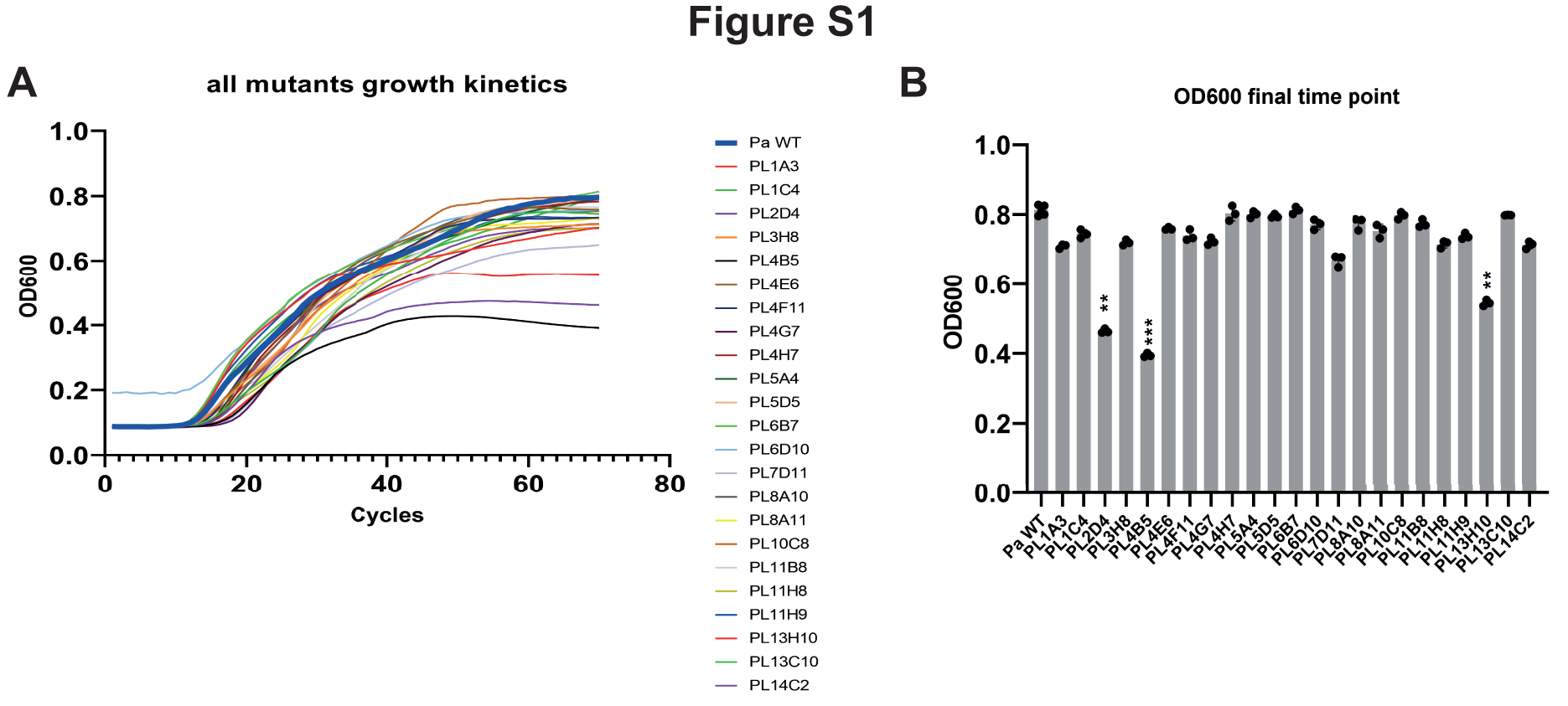

### Supplemental figure 2

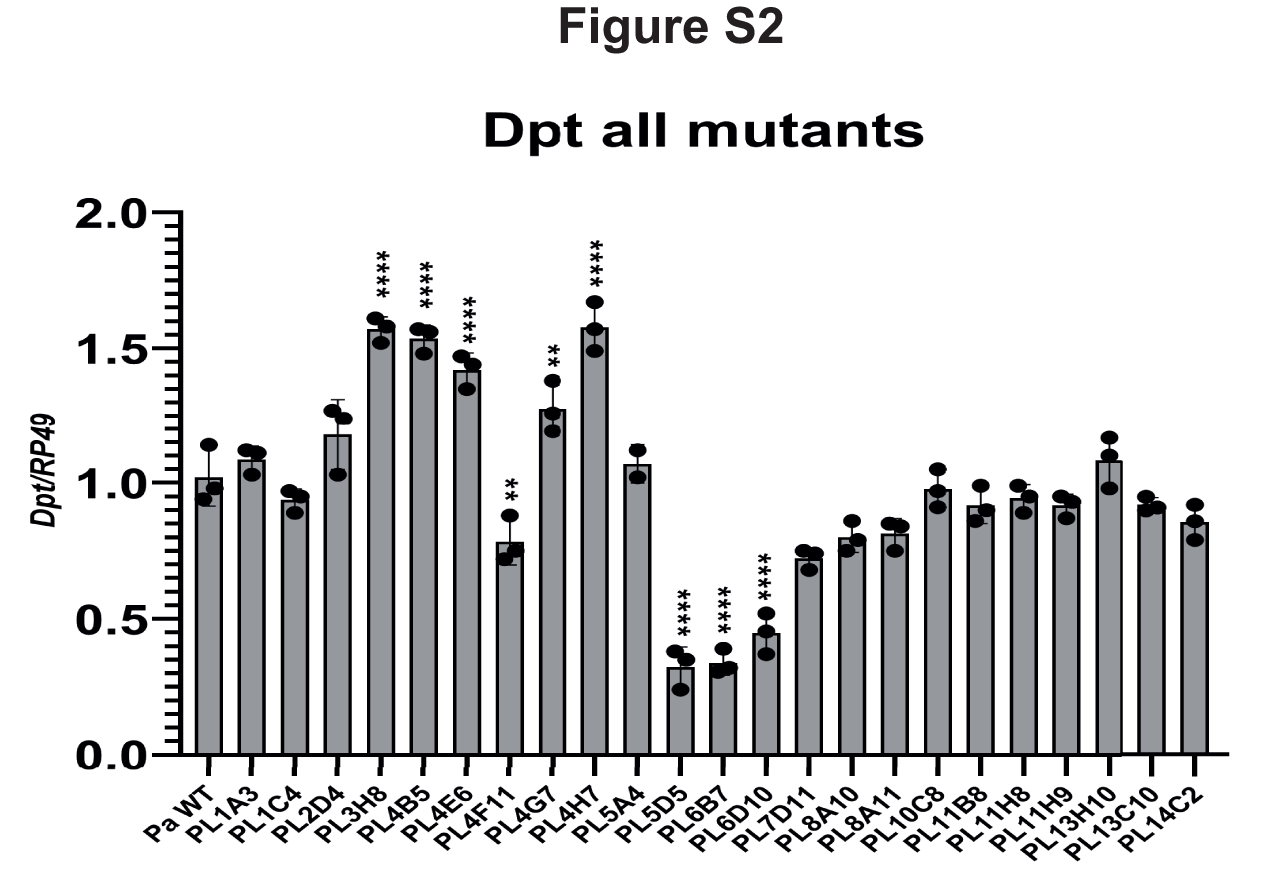

### Supplemental figure 3

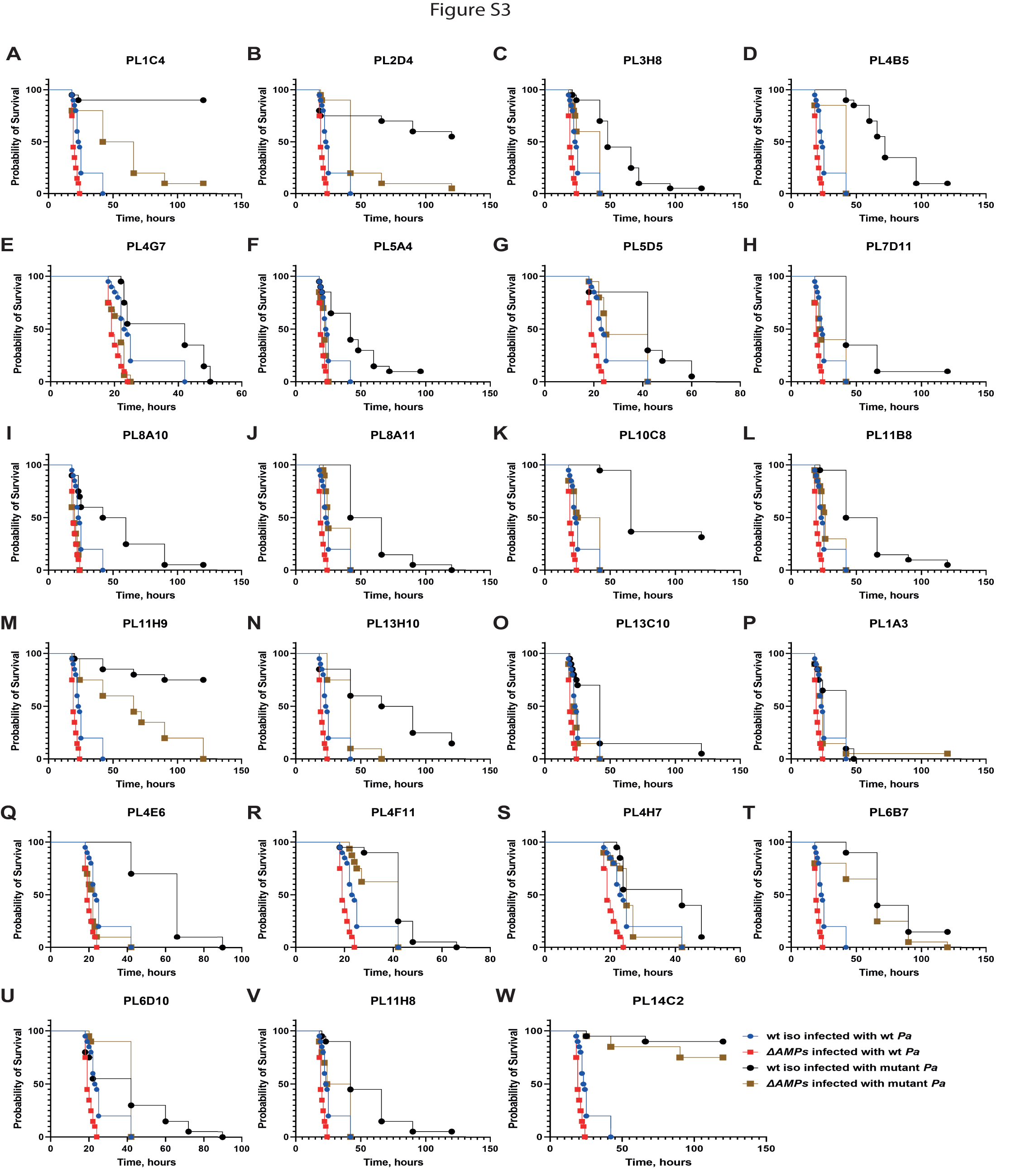

### Supplemental figure 4

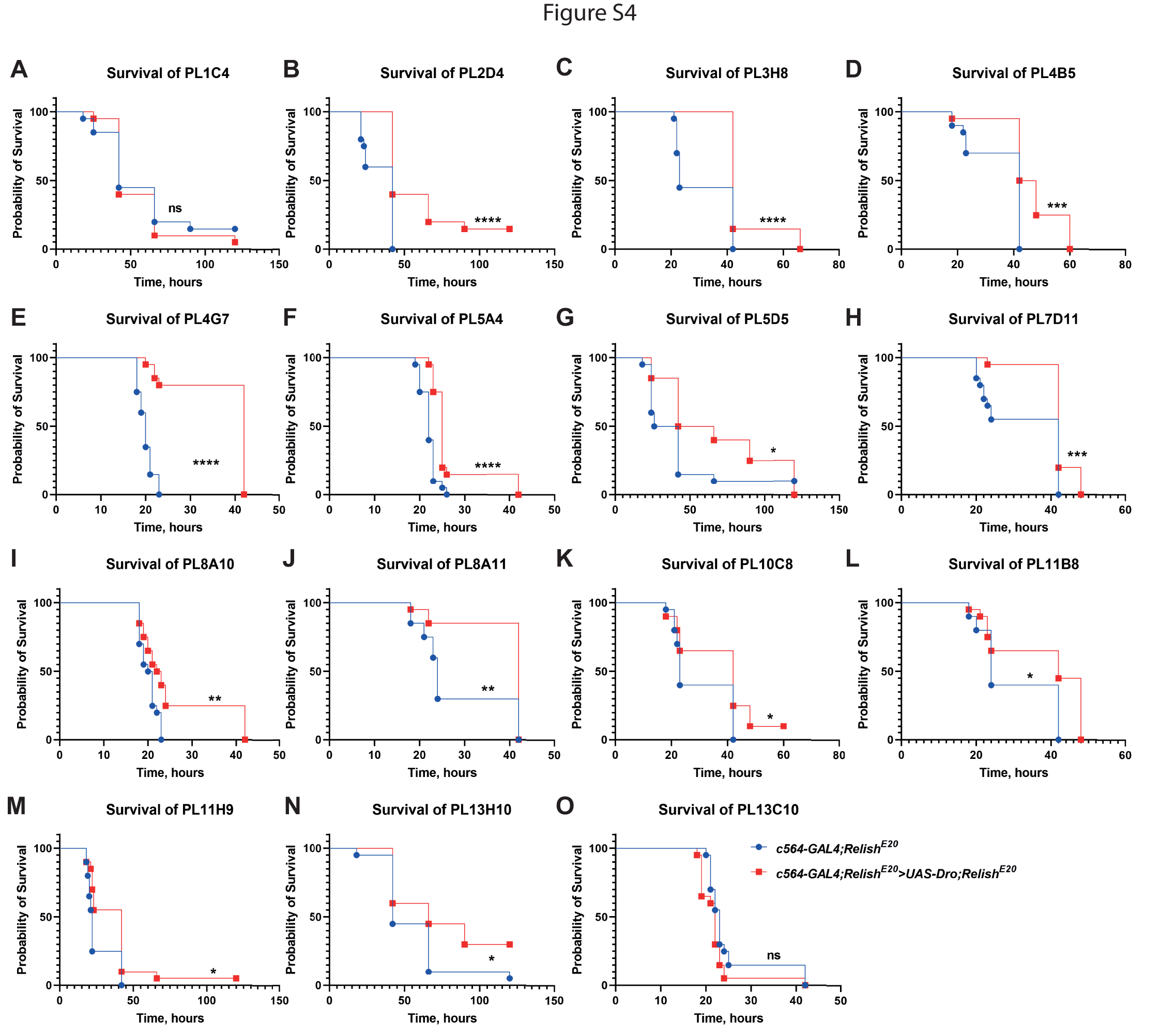

### Supplemental figure 5

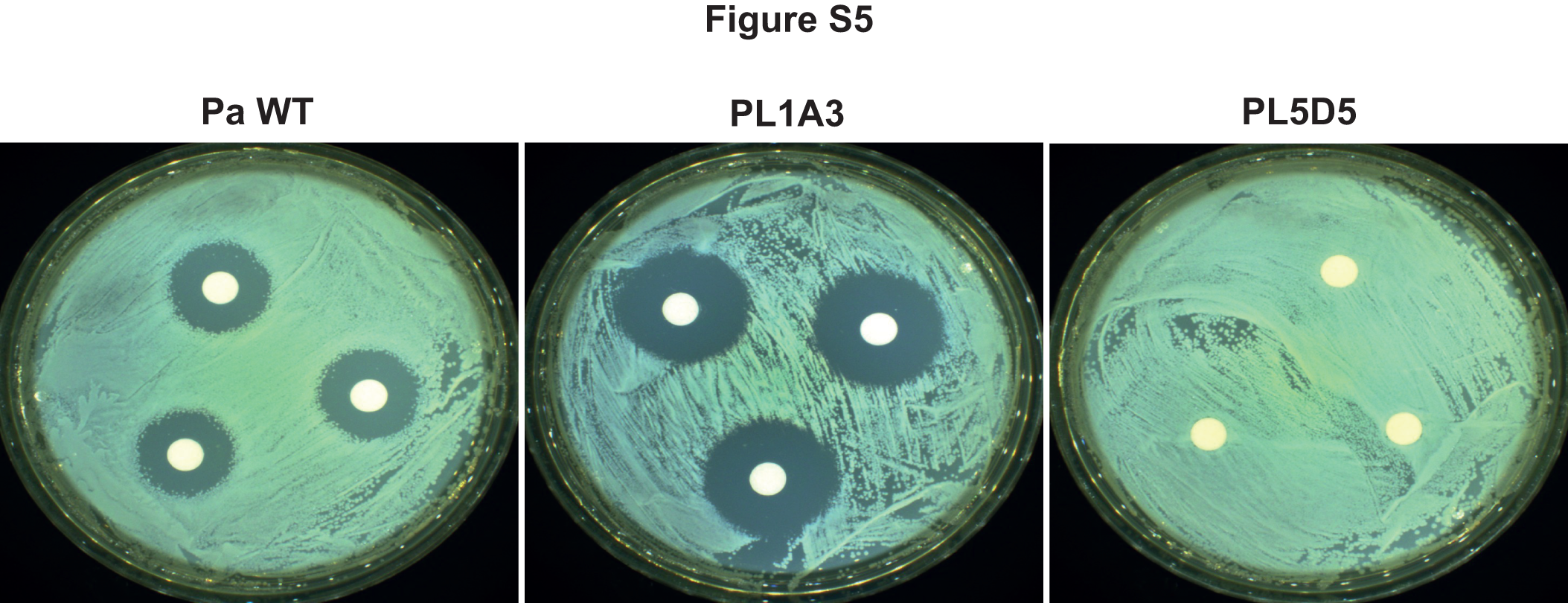

### Supplemental figure 6

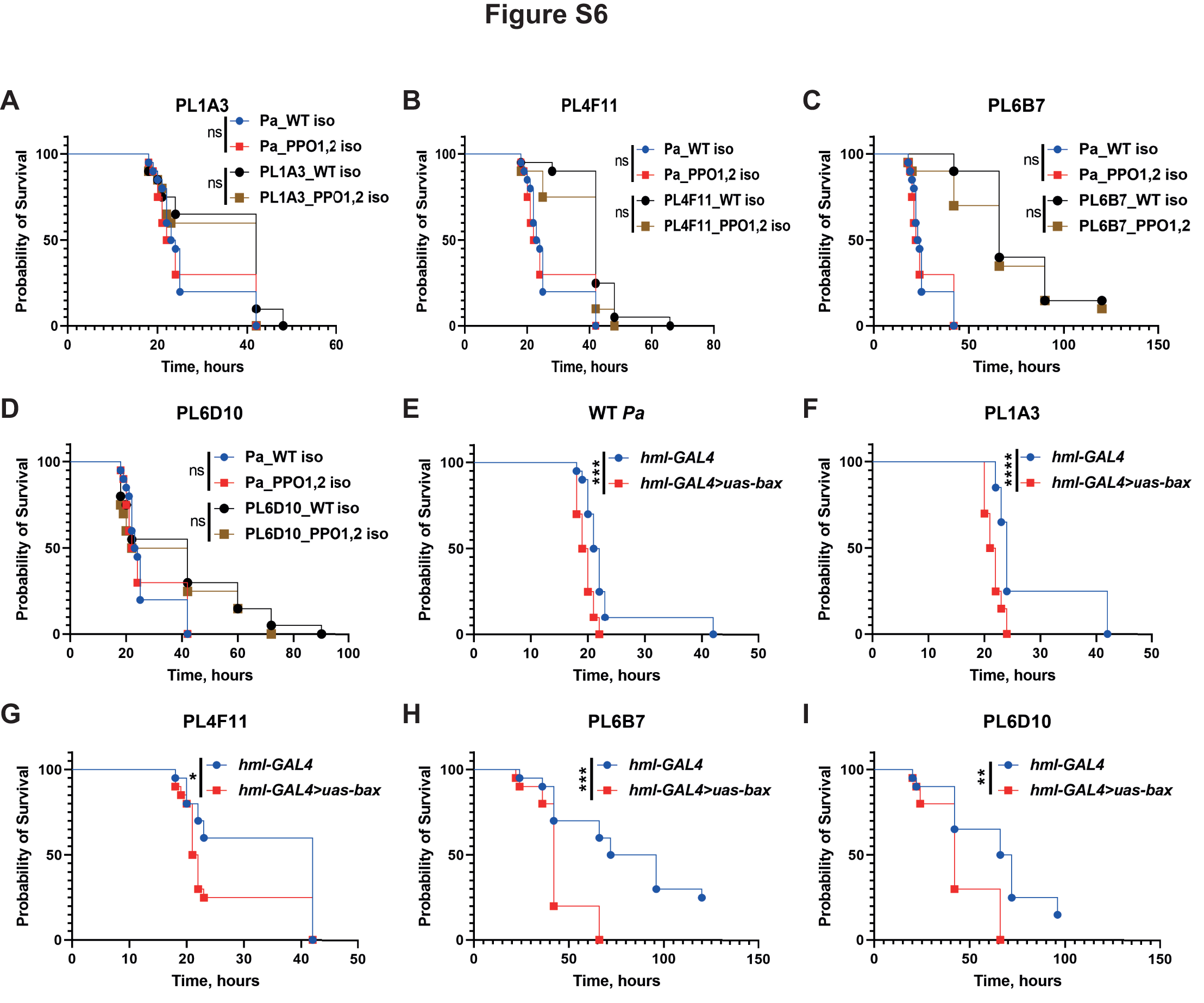
